# Necrosis reduction efficacy of subdermal biomaterial mediated oxygen delivery in ischemic skin flaps

**DOI:** 10.1101/2022.07.31.502155

**Authors:** Arghavan Rastinfard, Benjamin Dalisson, Mirko Gilardino, Kevin Watters, Dario Job, Geraldine Merle, Arturo Vela Lasagabaster, Yassine Ouhaddi, Jake Barralet

## Abstract

Inadequate tissue blood supply (e.g., in a wound or a poorly vascularised graft) can result in tissue ischemia and necrosis. As revascularization is a slow process relative to the proliferation of bacteria and the onset and spread of tissue necrosis, extensive tissue damage and loss can occur. Necrosis can spread rapidly, and treatment options are limited such that loss of tissue in ischemic tissue following necrosis onset is considered unavoidable and irreversible.

Oxygen delivery from biomaterials exploiting aqueous decomposition of peroxy-compounds has shown some potential in overcoming the supply limitations caused by quite short oxygen diffusion distances in tissues by creating higher concentration gradients than can be attained by air saturated solutions or by distributing oxygen supply throughout a scaffold or construct by using particulate formulations. These have found application in tissue preservation, bioinks, creation of 3D tissue analogues etc. In preclinical models among the more exciting reports was a single study demonstrating reduction of ischemic skin necrosis albeit only short term using short term sub dermal delivery of oxygen below ischemic skin flaps. To explore this effect further, we developed an implantable solid peroxide-biomaterial based system with reduced hydrogen peroxide release by virtue of incorporation of minerals to catalytically decompose it in a much longer flap than examined previously. Blood flow in this flap reduced from essentially normal to essentially zero, along its 9cm length. Without treatment ∼50% of the total flap was necrotic in 2-4 days. In both groups, complete necrosis in the distal third of the flap with no observable flood flow was observed. But in the middle low blood flow region of the flap, treatment did prevent necrosis. This study indicated that subdermal oxygen delivery alone cannot completely mitigate dermal necrosis if no blood flow is present, but it could improve the survival of partially tissue at least in the short term which could find application to augment conventional treatments or to gain time until surgical intervention.

## 1. Introduction

The pedicelled skin flap is an important surgical approach to skin reconstruction. When the flap is created, part of the vascularization is cut, and if its length exceeds that supplied by a single angiosome, the resulting blood flow path becomes random and is effected by choke vessels that ‘randomly’ connect discrete otherwise unsupplied angiosomes. These flaps are referred to as random flaps, and the ability of the skin of these flaps to survive becomes a function of the length of the pedicle [1]. In addition, comorbidities like diabetes, radiotherapy, etc., may further exacerbate the ability of the blood supply to remodel sufficiently rapidly to prevent tissue death. Ischemia, a restriction of blood flow to tissues, causes a shortage of oxygen and nutrients that are needed for cellular metabolism [2] and results in a loss of tissue homeostasis and can result in damage or dysfunction of tissue when delivery fails to meet metabolic requirements. In blood, the glucose concentration is regulated between 1.4 mmol/L and 6.2 mmol/L [3] and oxygen between ∼100 to 150 µmol/L [4]. Without oxygen, the adenosine triphosphate (ATP) production cycle is limited to a glycolytic yield of 2 moles of ATP for 1 mole of glucose. In the absence of oxidative phosphorylation, glucose is converted into lactate through glycolysis, and the intracellular pH decreases [2, 5]. The lack of ATP leads to intracellular Na^+^, water, and Ca^2+^ accumulation, cell membrane depolarization, protease activation as well as an increased reactive oxygen species (ROS) production [6], all of which can irreversibly damage cells and lead to necrosis. In addition, in this environment, the mitochondrial membrane is disrupted and opening the permeability transition pore, which further decreases the ATP production and releases apoptotic factors, and initiates the apoptotic cascade [2, 7]. These alterations and thus the degree of tissue injury varies with the extent and duration of the ischemic period [2]. In addition, necrosis can spread to surrounding tissue, and treatment options are still currently so limited that loss of tissue in ischemic limbs or wounds is considered unavoidable [8]. However, while necrotic tissue is considered an impediment to healing being sloughed off or debrided, many ‘dead’ and deliberately cell-free acellular tissues are used as regenerative allogenic scaffolds [9-11]. This then suggests, that with modification, necrotic tissue could theoretically actually support the healing process.

Calcium peroxide can store and release oxygen upon exposure to aqueous conditions, decomposing to form molecular oxygen and hydrogen peroxide [12, 13]. This and other peroxy-compounds have been widely studied for various oxygen delivery applications, for example, remediating waterlogged soil and in aquaculture [14-17], and tooth whitening [18-20], but in implantable tissue repair applications, they are still at a research level and have not found clinical application. The main reasons for this are that hydrogen peroxide may also be formed and that caustic hydroxides may be formed. We previously determined that toxic effects could be attributed to hydrogen peroxide generation [12] and pH elevation and so attempted to mitigate this by adding iron oxide as a peroxide catalyst, adding an acidic buffer to help maintain neutrality, and by reducing the rate of reaction with a polymer diffusion barrier. Since we previously determined that at high pH (as found at the particle surface), carbonate ions can reduce hydrogen peroxide levels [12], we offset the acidity of calcium monophosphate by adding sodium bicarbonate. This study aimed to evaluate the preclinical efficacy of this oxygen generating subdermal film to reduce ischemic skin necrosis in full thickness rat random skin flaps.

## 2. Materials and Methods

### 2.1. Fabrication of subdermal oxygen generating films

Calcium peroxide (CaO_2_, FMC Corporation), iron (II, III) oxide (Fe_3_O_4_, Sigma-Aldrich), sodium bicarbonate (NaHCO_3_, Sigma-Aldrich), and calcium dihydrogen phosphate monohydrate, (Ca(H_2_PO_4_)_2_.H_2_O, abcr GmbH) powders were sieved using 74 µm sieve (ASTM 200 mesh) and then mixed. To fabricate subdermal oxygen releasing films, 10% PCL solution (w/v, M_w_ 80000, Sigma-Aldrich) was prepared by dissolving PCL in chloroform (Fischer Scientific). The particle mixtures were added to PCL solution (Table 1) and stirred for 24 hours to provide homogenous distribution of particles.

**Table 1.** Weight percentage (%wt) of the components composing experimental and control groups.

| Sample | Weight percentage (% wt) of the components |  |  |  |  | Total |
| --- | --- | --- | --- | --- | --- | --- |
| | $\text{CaO}_2$ | $\text{Fe}_3\text{O}_4$ | $\text{NaHCO}_3$ | $\text{Ca}(\text{H}_2\text{PO}_4)_2 \cdot \text{H}_2\text{O}$ | Glass beads | |
| Experimental | 13.7 | 13.7 | 3.3 | 2.7 | 0 | 33.4 |
| Control | 13.7 | 0 | 0 | 0 | 19.7 | 33.4 |

In order to produce a control for in vitro measurements that lacked buffering and catalyst components, a suspension of 13.7% w/w CaO_2_ and 19.7% w/w glass beads (212-300 µm, Sigma-Aldrich) were prepared in a 10%w/v PCL in chloroform. The glass beads acted as a filler so that the amount of particulates in both experimental and control groups was similar, since it is well known that polymer permeability to water is strongly affected by filler level [21-23]. They were also sieved through a 74 µm sieve, same as buffering and peroxidase components to approximate the volume of the experimental sample filler. The composite solutions were cast in crystallizing dish, and the solvent was evaporated overnight.

2.3g of cast films were used to prepare the implants for *in vitro* oxygen release measurements and *in vivo* studies. PCL films become malleable around melting point (60 °C), so the dry films were molded into 1.7×7×0.1 cm^3^ implants by heating them on a laboratory hot plate.

### 2.2. pH value and hydrogen peroxide measurements

pH and hydrogen peroxide (H_2_O_2_) measurements were carried out in a sealed chamber for 14 days. Experimental and control samples containing 0.7 mg CaO_2_ were immersed in 2 mL of water. The amount of released H_2_O_2_ was measured by the Thermo Scientific peroxide assay kit. The pH value of the solution was also measured by a pH meter (Mettler Toledo InLab Microelectrode).

### 2.3. Cytotoxicity of subdermal oxygen generating films

The samples were sterilized in 70% ethanol for 1 hour, washed three times with PBS, and dried under tissue culture hood overnight. 24-well plate and cell culture insert with 0.4µM pore size (PET track-etched membrane, Corning Inc, Corning, NY, USA) were used for cell culture studies. Madin-Darby Canine Kidney (MDCK) cells were cultured in RPMI-1640 (R8758, Sigma-Aldrich) supplemented with 1% penicillin/streptomycin and 10% fetal bovine serum. MDCK cells were seeded at a seeding density of 2 × 10^4^ cells/well and kept overnight at 37°C and 5% CO_2_. Subsequently, each well medium was replaced with a fresh medium (700 µL). Oxygen generating films with the same amount of CaO_2_ (0.17 mg) were each placed individually in culture inserts of a 24 well plate, and then 300 µL of the medium was added. No material addition cultures were used as a positive control. After 24 hours, the cell viability was examined.

### 2.4. Live-dead assay

Cell viability was examined using a live-dead assay (Molecular Probes TM, LIVE/DEAD Viability/Cytotoxicity Kit). A dye solution containing 2µM Calcein AM and 2µM Ethidium homodimer-1 was prepared. After 24 hours, the cell culture inserts and medium were removed, and 300µL of dye solution was added to each well and then incubated for 30 min in the incubator. Cells after staining were analyzed by fluorescence microscope (EVOS M5000 Imaging System).

### 2.5. Oxygen and lactate measurements

For *in vitro* measurements, an oxygen generating film was immersed in 40mL phosphate-buffered saline (PBS) at 25°C in an open beaker (Ø=5cm, 2cm liquid depth). The oxygen content of the PBS was assessed using an oxygen probe (AL300 Oxygen Sensor Probe, Ocean Optics). For calibration, the oxygen sensor was exposed to 0, 10, 20, 30, 40, 60, and 80% dissolved oxygen in water (prepared by flushing nitrogen and oxygen gas through water) until the Tau value (fluorescence lifetime) stabilize. For *in vivo* measurements, the probe was inserted through the gaps between sutures under the skin flap at 3 different positions on the flap: proximal, middle, and distal. Measurements were performed under anesthesia just after the surgery (day 0) and on days 1, 2, 4, 6, 8, 10. At the moment of euthanasia, sections of the flap were frozen at - 80°C and processed [24] to measure the lactate content of the tissues using a Lactate assay kit (Sigma-Aldrich).

### 2.6. Surgical methods

Wistar rats (male, 5 to 6-months old, 500 to 600g, Charles River Laboratories Inc. Montreal, QC, Canada) were randomized into 2 groups. The control group received no biomaterial, and the experimental groups received oxygen releasing films subdermally (N=9 per group). Surgical procedures were performed after approval from McGill University Animal Care Committee (UACC, #7899). Animals received carprofen (10mg/kg) 30 min prior to the surgery; all surgeries were performed under general anesthesia using 2% isoflurane. Full depth skin flaps of 9 × 2 cm^2^ in size were created on the back [25]. A silicon sheet (SMI, 0.005” NRV M/M 40D) was placed over the muscle to prevent revascularization and reperfusion of the dermal side of the flap from the underlying tissue, then the film was positioned distally, and the skin was re-replaced on top of it and sutured. Animals received carprofen (10mg/kg) every 24 hours for 3 days post-surgery, then slow-release buprenorphine every three days until the experimental endpoint. Animals were allowed free access to food and water and housed in a 12 h day/night cycle. Five animals were sacrificed on day 6, four animals on day 10 using carbon dioxide.

### 2.7. Evaluation of skin flap survival

The skin flap areas were photographed on days 1, 2, 4, 6, 8, and 10 after surgery. On each day, ImageJ software was used to analyze the images and measure the necrotic area as determined by black discolouration of the skin. The necrotic tissue was calculated by this formula: necrosis area/initial flap area × 100.

### 2.8. Histology and immunohistochemistry analysis

Flaps sections (proximal, middle, and distal) were cut and divided in half lengthwise. A half sections were used for lactate measurement, and other sections were fixed in 4% paraformaldehyde for 24 hours at 4°C. Samples were placed in a tissue Tek VIP5 Vacuum Infiltration Processor and embedded in paraffin by the Leica EG 1150H. Then, tissues were sectioned with a microtome (RM 2255 microtome Leica Microsystems, Germany) in 4µm of thickness size. Tissue sections were deparaffinized using xylene and stained with Hematoxylin & Eosin (H&E). Immunohistochemical staining was performed with anti-CD34 for blood vessels (1:1000 dilution, ab81289, Abcam), anti-Foxp3 antibody for regulatory T cells (1:200 dilution, ab22510, Abcam), anti-iNOS antibody for M1 polarized macrophages (1:100 dilution, ab3523, Abcam), recombinant anti-Liver arginase antibody for M2 polarized macrophages (1:1000 dilution, ab203490, Abcam), and anti-F4/80 as general macrophage marker (1:500 dilution, EMR1 Antibody (PA5-21399)) using standard procedures. Each slide was washed with PBS (pH 7.4) and incubated with a secondary antibody using the Vectastain ABC kit (Vector Laboratories) according to the manufacturer’s instructions. The color was developed with 3,3’-diaminobenzidine (DAB) solution (Vector Laboratories) containing hydrogen peroxide. The sections were scanned using Aperio scanner ScanScope XT and analyzed by ImageScope software (Leica Biosystems). Tissue necrosis area visible in H&E-stained sections was measured. For microvascular and macrophage markers, DAB staining was quantified within the distal, middle, and proximal areas using the Color Deconvolution algorithm (Aperio). The results were reported as percent total positive × total stained area/total analysis area.

## 3. Results

### 3.1. *In vitro* characterization of subdermal oxygen generating films

First, we examined the buffering ability of oxygen releasing films in offsetting pH values. When immersed in water, the addition of the control raised the pH to 10.8 within 2 hours and then continued to increase to 11.4 and 11.6 over 2 and 14 days. In contrast, oxygen releasing films increased pH value to 8.1 within 2 hours and pH remained below 10 over 2 days. Then, pH increased gradually to 11.6 within 2 weeks (Fig.1A). These measurements were initially conducted in Eppendorf tubes that were closed in between measurements. Repeating the experiment in tubes that were left open resulted in∼20% lower pH for both experimental and control groups due to carbonation with atmospheric CO_2_. The pH value for the control group peaked at 10.1 after 2 hours and then decreased to 8.7 over 24 hours. In contrast, oxygen generating group caused a rise in pH to 8.4 and then stabilized and reached a plateau of 8.7 pH after 24 hours (Fig. 1B).

**Fig. 1.**
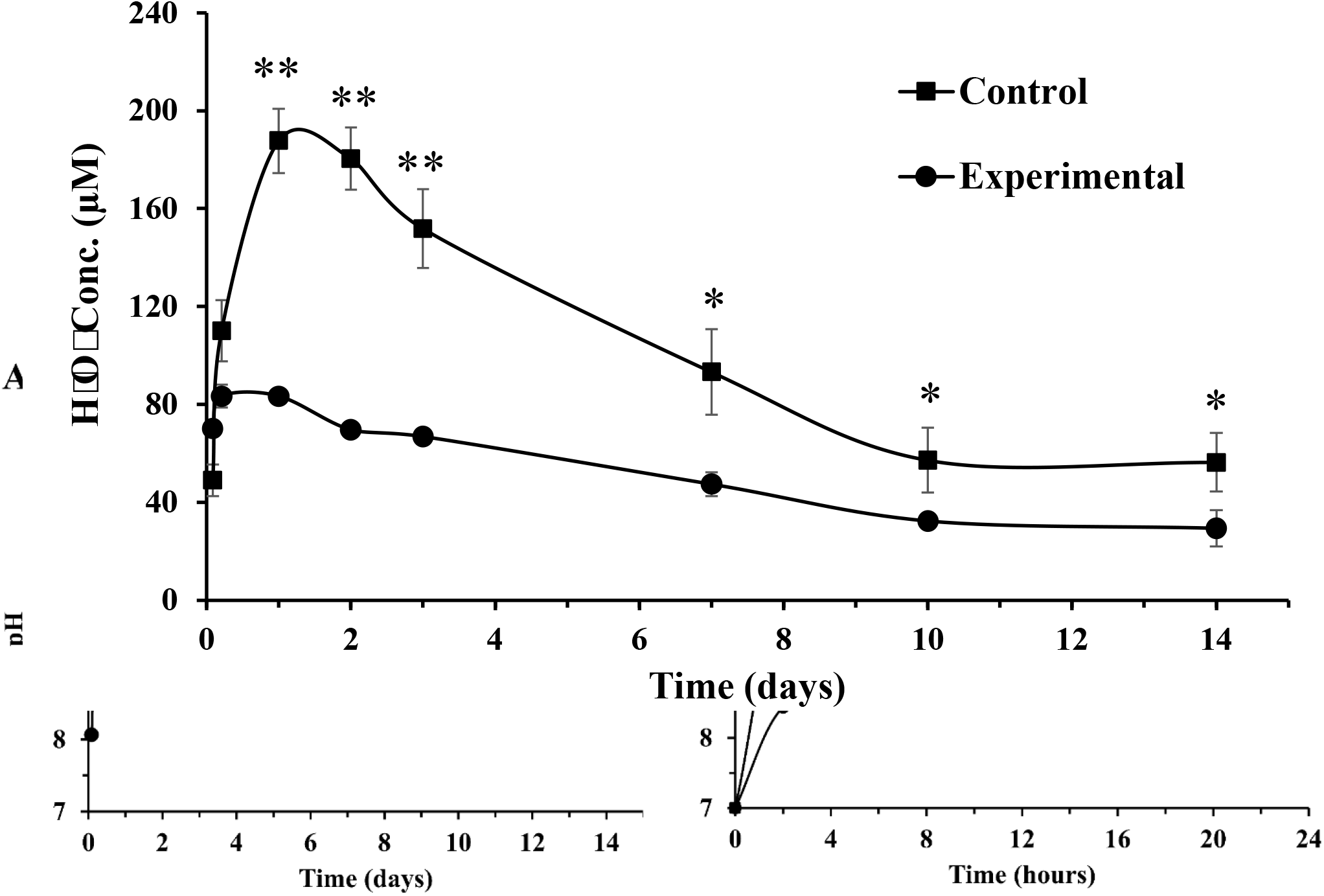
The pH value of water containing control and experimental films (a) over two weeks in Eppendorf tubes without air flow (b) over 24 hours in open Eppendorf tubes with air contact. The control group is oxygen releasing films formed without iron oxide nanoparticles and buffering components. N=3.

We also attempted to reduce the potentially cytotoxic effect of hydrogen peroxide through incorporating iron oxide with known catalytic activity towards hydrogen peroxide decomposition [26, 27] and sodium bicarbonate since we previously found that carbonate could also be effective in reducing hydrogen peroxide levels [12]. Over the course of the experiment (Fig. 2), the experimental and control groups reached the maximal hydrogen peroxide release after one day, 83.3µM, and 187.7µM for experimental and control, respectively, (P<0.05). Hydrogen peroxide content decreased gradually for both groups, indicating that decomposition rate exceeded the release rate.

**Fig. 2.** The analysis of hydrogen peroxide release from control and experimental films over two weeks in closed Eppendorf tubes. The control group is oxygen releasing films formed without iron oxide nanoparticles and buffering components. N=4, **P<0.01 and *P<0.05.

Next, we examined the cytotoxicity of experimental and control on MDCK cells after 24 hours. As shown in Fig. 3, the control group is significantly cytotoxic with only 34% cell viability. In contrast, the viability if cells in the experimental group was 81%. Therefore, oxygen generating films incorporating iron oxide and buffering components enhanced cytocompatibility since iron oxide reduced hydrogen peroxide release and buffers controlled shifting pH values.

**Fig. 3.**
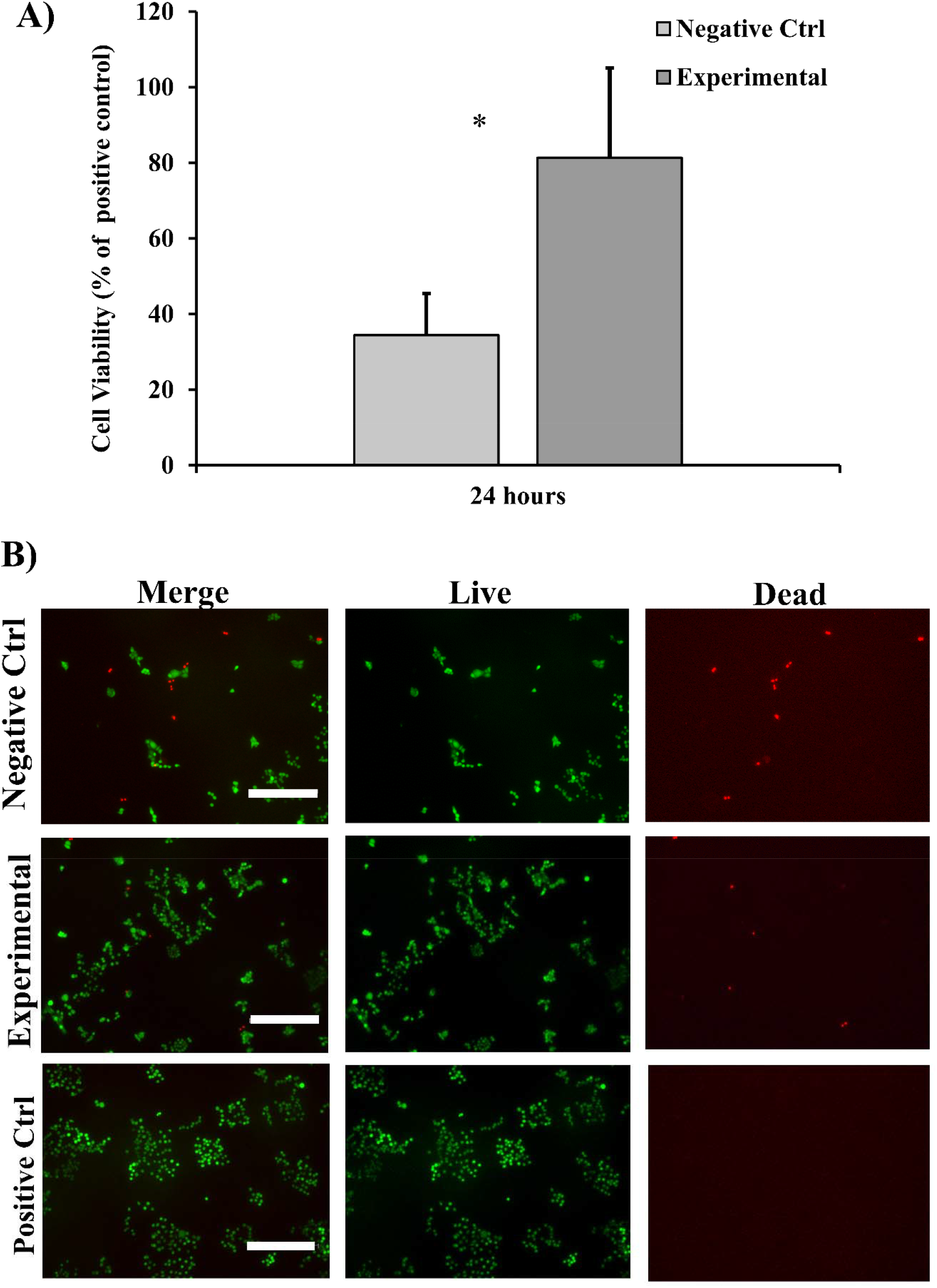
(A) Cytotoxicity of experimental and control group (B) Fluorescent microscopic images of the cells after 24 hours using live/dead assay. N=3, * P<0.05. The negative control group is oxygen generating films formed without iron oxide nanoparticles and buffering components. Positive control is cells without any treatment. Scale bar is 300µm.

The oxygen release from experimental was significantly higher compared to that from control baseline remained at about 20% throughout the entire experiment (P<0.05), Fig. 4. There was a burst release of oxygen at first 15 minutes and oxygen generating films delivered about 35% oxygen. After that, oxygen levels raised to about 48% in day 1 and reached the maximum amount of 52% on day 2. Although oxygen levels gradually decreased after day 2, oxygen generating film was able to sustain oxygen release over two weeks.

**Fig. 4.**
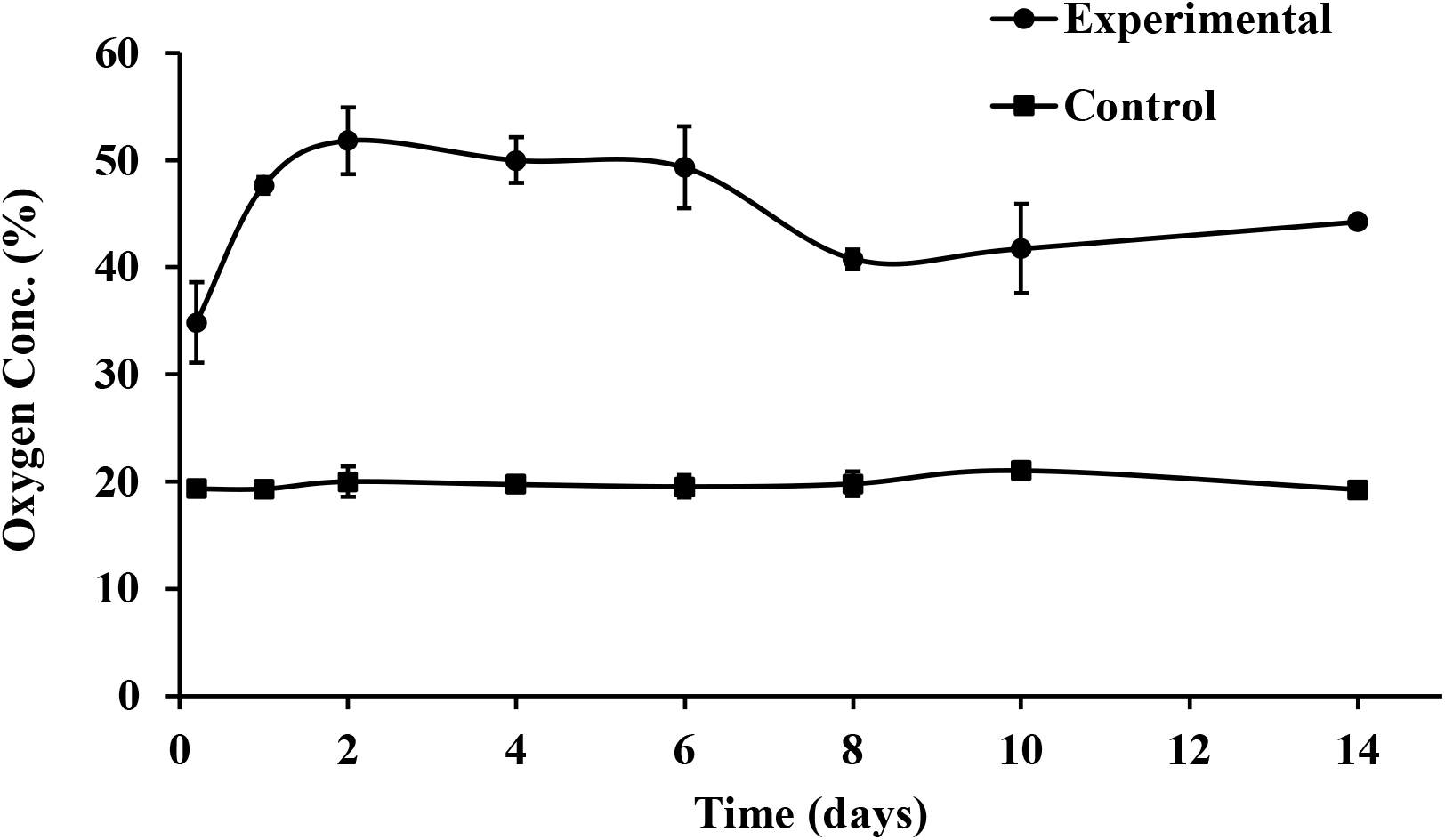
Oxygen release from experimental and control groups into 40 mL PBS in containers open to air at room temperature for two weeks.

### 3.2. *In vivo* characterization

At each time point, the skin flaps were photographed, and the necrotic area was evaluated. Immediately after the surgery, the distal part of the flaps of both groups appeared slightly blue (Fig. 5A) and dark blue after one day with some necrosis at the extremity. On day 2, the necrosis was obvious in the distal portion of the flap for both groups. The middle and proximal portions were normal in the subdermal oxygen generating group, while in the control group only proximal area appeared to be in the normal range. On day 4, the dark blue middle portion in control group turned brown and gradually blackened over the next few days. The necrotic area in the oxygen releasing group became darker and rougher, starting day 4, but the necrosis did not spread to the middle and proximal area over the experiment.

**Fig. 5.**
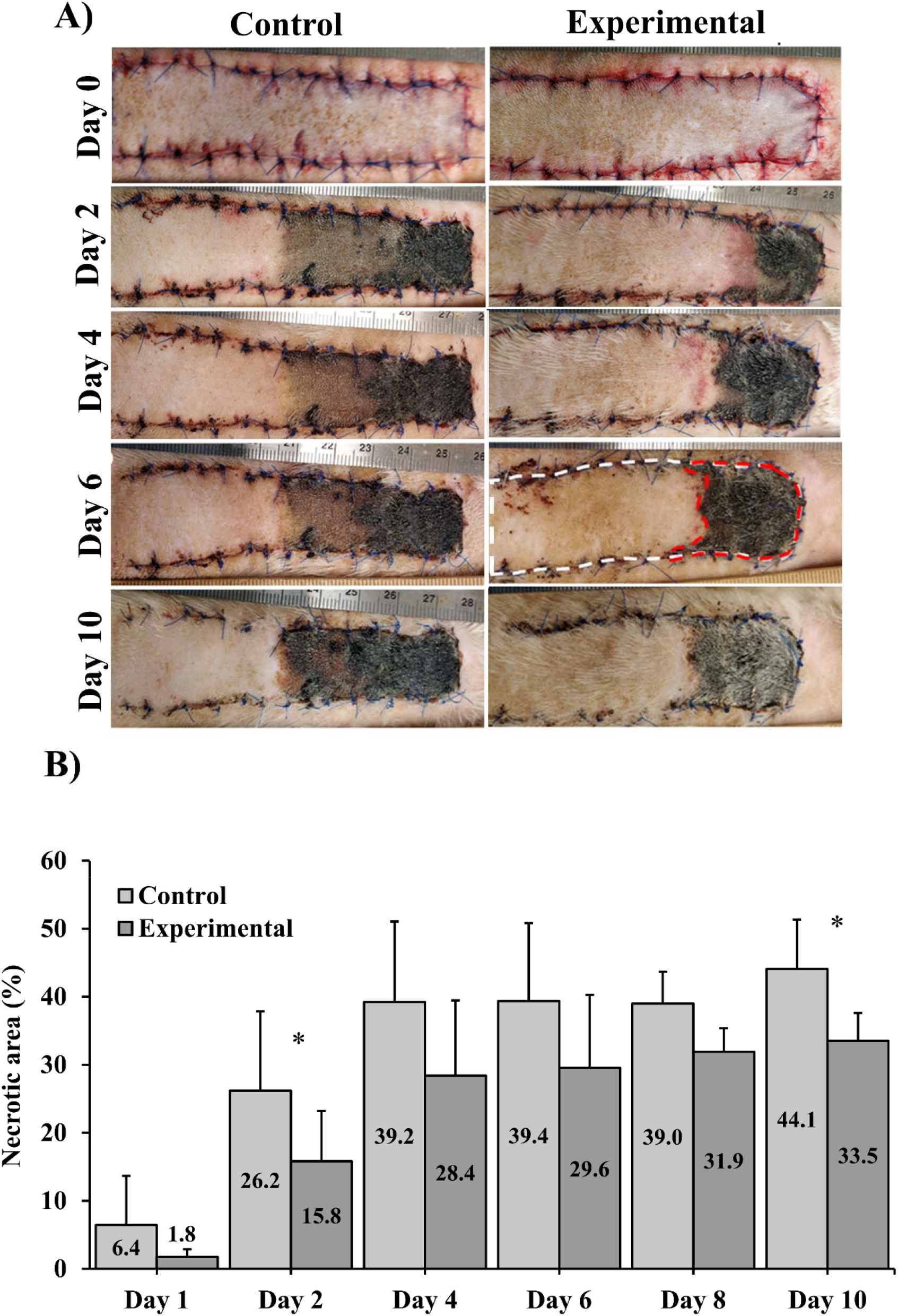
(A) Representative photographs of the skin flap on days 0, 2, 4, 6, and 10 for control and experimental groups, (B) Histogram representing the visible relative necrotic area over time, expressed as necrotic area (red dotted line) over total visible flap area (white dotted line). Control group is skin flaps without receiving any materials. Values are expressed as mean±SD, N=9 per group. *P<0.05.

The area of skin flap necrosis was significantly greater (P<0.05) in the control group than in the experimental group on days 2 and 10 (Fig. 5B). In the control group, the necrotic surface area significantly increased after surgery and remained at around 40±10% from day 4 to day 10. In the experimental group, the necrosis increased from about 2±1% on day 1 to 28±11% on day 4 and then remained almost stable at 31±8% until the end of the experiment. Examination of the data revealed that animals developed necrosis at very different rates even within study groups, using analysis of a dichotomous endpoint (Did extent of necrosis exceed 40% at any time during healing in a particular subject? (Y/N)), we determined a statistical difference of P<0.05 at a power of 98.4% (Table 2).

**Table 2.** Result of dichotomous endpoint analysis.

| Sample | Size | Proportion of Necrosis > 40% | Results |
| --- | --- | --- | --- |
| Experimental | 9 | 11% | P <0.05 |
| Control | 9 | 89% | Post-hoc power 98.4% |

Subcutaneous oxygen concentration was measured immediately after the surgery and on days 1, 2, 4, 6, 8, and 10 post-surgeries in the middle of the proximal, middle and distal third sections. In the control group, no significant differences in subcutaneous oxygen level were observed from day 1 to day 10 at any sections (Fig. 6). A similar observation was recorded in the experimental group. Oxygen concentration normally found in tissues (physoxia) is between 2 and 5%, and oxygen levels less than 2% are considered hypoxic [28]. As shown in Fig. 6, the oxygen level in the experimental group at any sections of the flap was near physoxic concentration, while there was hypoxia in the control group. The subcutaneous oxygen concentration in the middle and proximal portions was found to be significantly higher in the subdermal oxygen releasing group than in control (P<0.001). In the distal portion, the oxygen level was not significantly different (P<0.05) between the experimental and control groups. To further investigate the impact of oxygen delivery on skin flap, we measured lactate levels of flap tissue. Higher lactate production can be an indication of glycolytic metabolism because of less oxygen content in the ischemic flap [24]. Fig. 7 showed no significant difference (P<0.05) in lactate concentration between experimental and control groups, although oxygen level was significantly greater in experimental than control. Moreover, lactate measurements for each section of the flap did not indicate significant differences in levels between flap regions or with time at days 6 and 10.

**Fig. 6.**
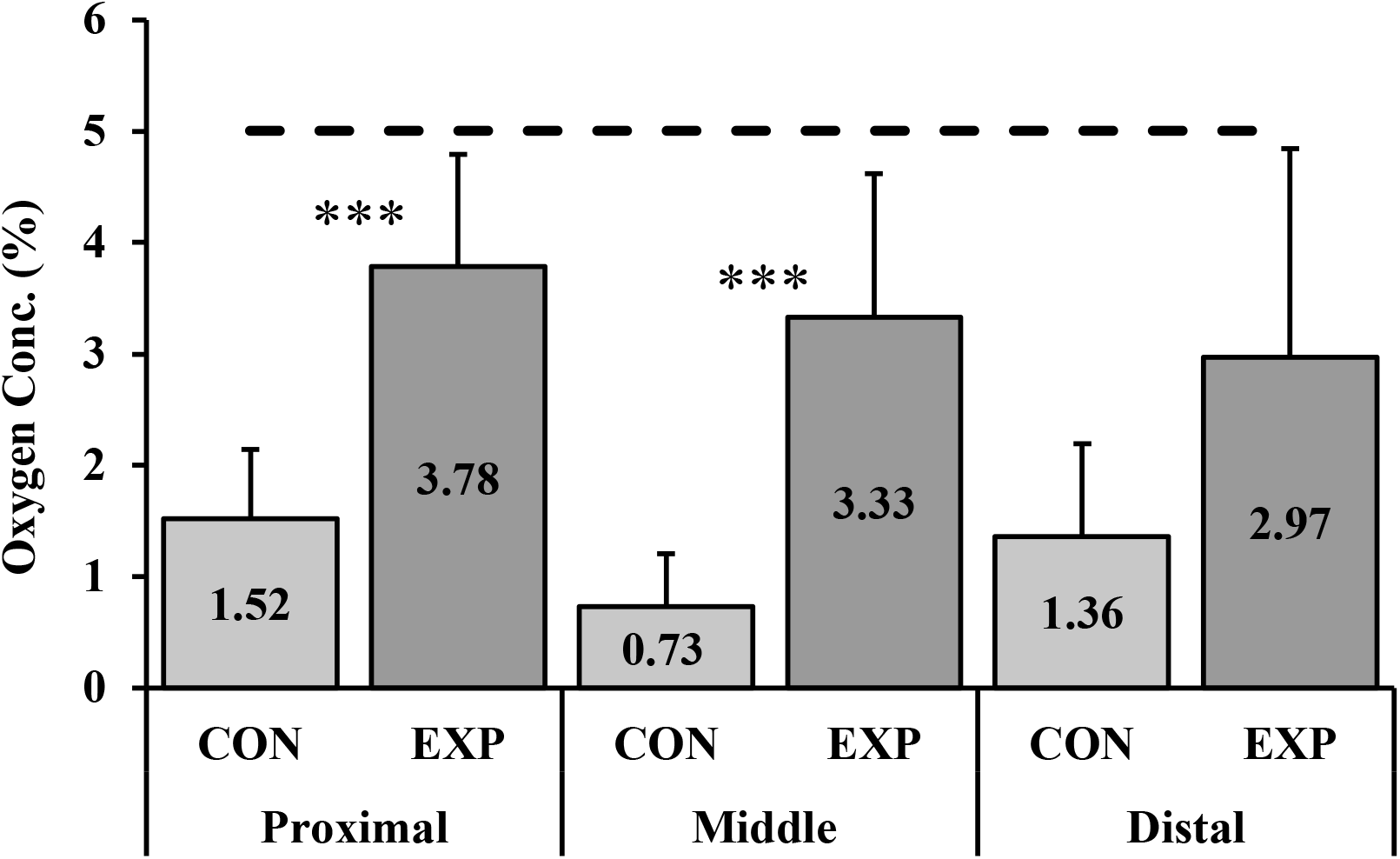
Histogram representing the subcutaneous oxygen concentration of the skin flaps under the proximal, mid-, and distal thirds of the flaps for both the untreated control and the experimental group, all time points combined (the dotted line represents physoxic oxygen concentration for comparison (2-5%)). Values are expressed as mean ± SD, N=8 per group. ***P<0.001. Abbreviations: CON: control, EXP: experimental.

**Fig. 7.**
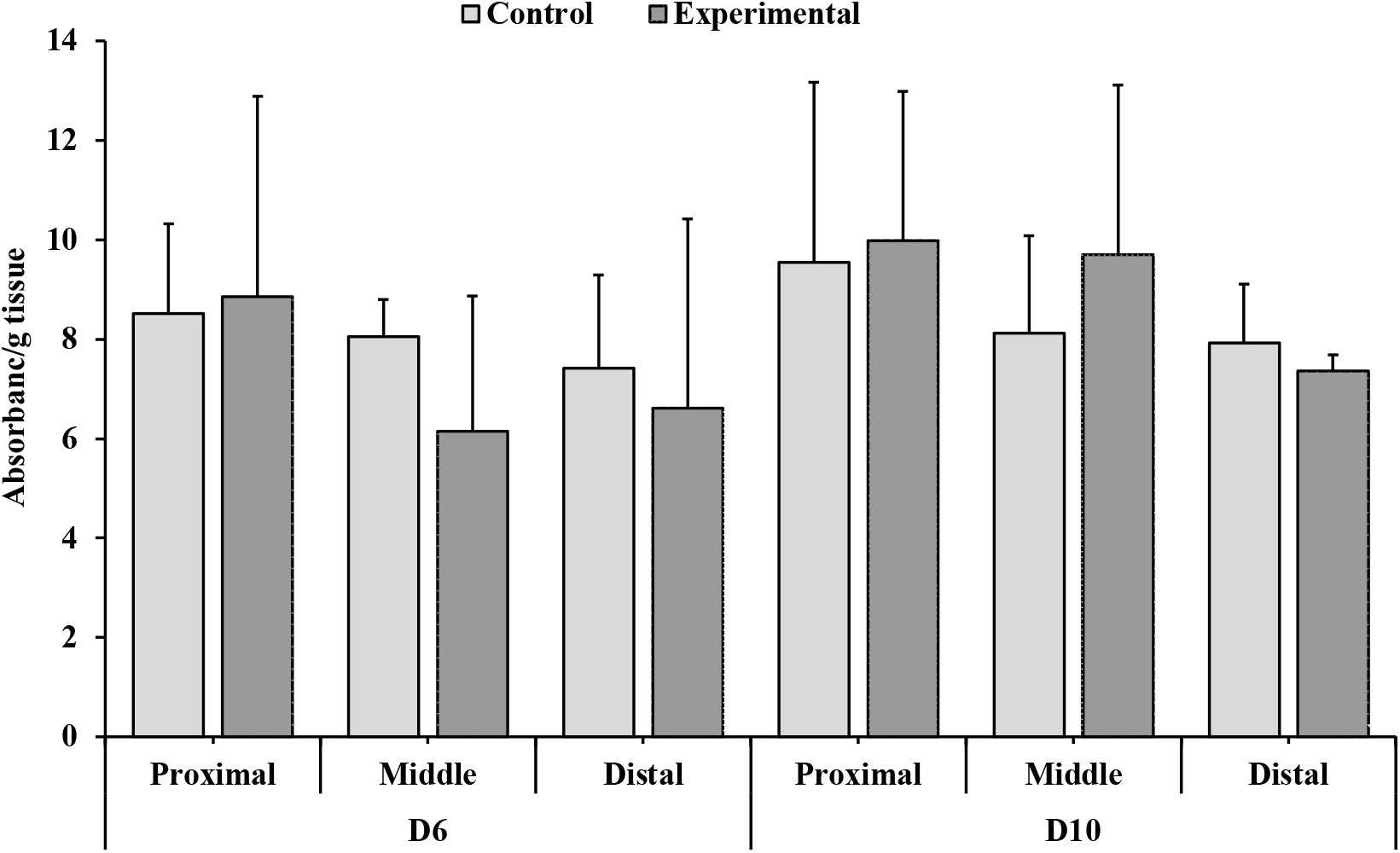
Histogram representing lactate quantification of each section of the flap for untreated control and experimental group at days 6 and 10. Results are expressed as absorbance per gram of tissue and were not significantly different between groups. Values are expressed as mean ± SD, N=3.

Fig. 8 shows an H&E-stained cross-section of skin flap tissues from both groups with two different magnifications. Little difference was visible in the tissue architecture of the proximal section when observed under H&E staining between experimental and untreated control groups (Fig. 8A, B). Fewer nuclei were discernible (indicative of necrotic non-viable tissue) for both groups as the distance from the proximal section increased. In the midsections, the experimental group exhibited necrosis to a lesser extent than controls, so the interface between live and dead tissue was located at the middle zone for control and distal or middle for the experimental group. Histologically, polymorphonuclear neutrophils (PMN) and also the formation of new epidermis underneath the necrotic tissue were visible at the interface between the necrotic and healthy tissues (Fig. 9). The distal portion was the necrotic area for both groups, where disruption of tissue architecture and increased eosinophilia were obvious. In the necrotic portion of the flap, ghost cells were visible around the appendages identified as preserved cell outlines without nuclei. For both groups, the distal sections mostly consisted of necrosed tissues resembling some of the aspects of coagulative necrosis [29, 30] with the disappearance of the nuclei and appendages. Histomorphometric measurement of the relative necrotic area in the skin flaps (epidermis, dermis, and hypodermis) was performed on the H&E-stained sections (Fig. 8C). On day 6, the mean necrotic area in the proximal third was 0.0% for both groups and in the distal sections 100.0 ± 0.0% and 85.5 ± 28.5% for the control and experimental group, respectively. In the middle section, the mean necrotic area was 49.3 ± 37.8% in the control group and 18.9 ± 9.8% for the experimental. There was no significant difference in necrotic area of the three sections between control and oxygen releasing groups.

**Fig. 8.**
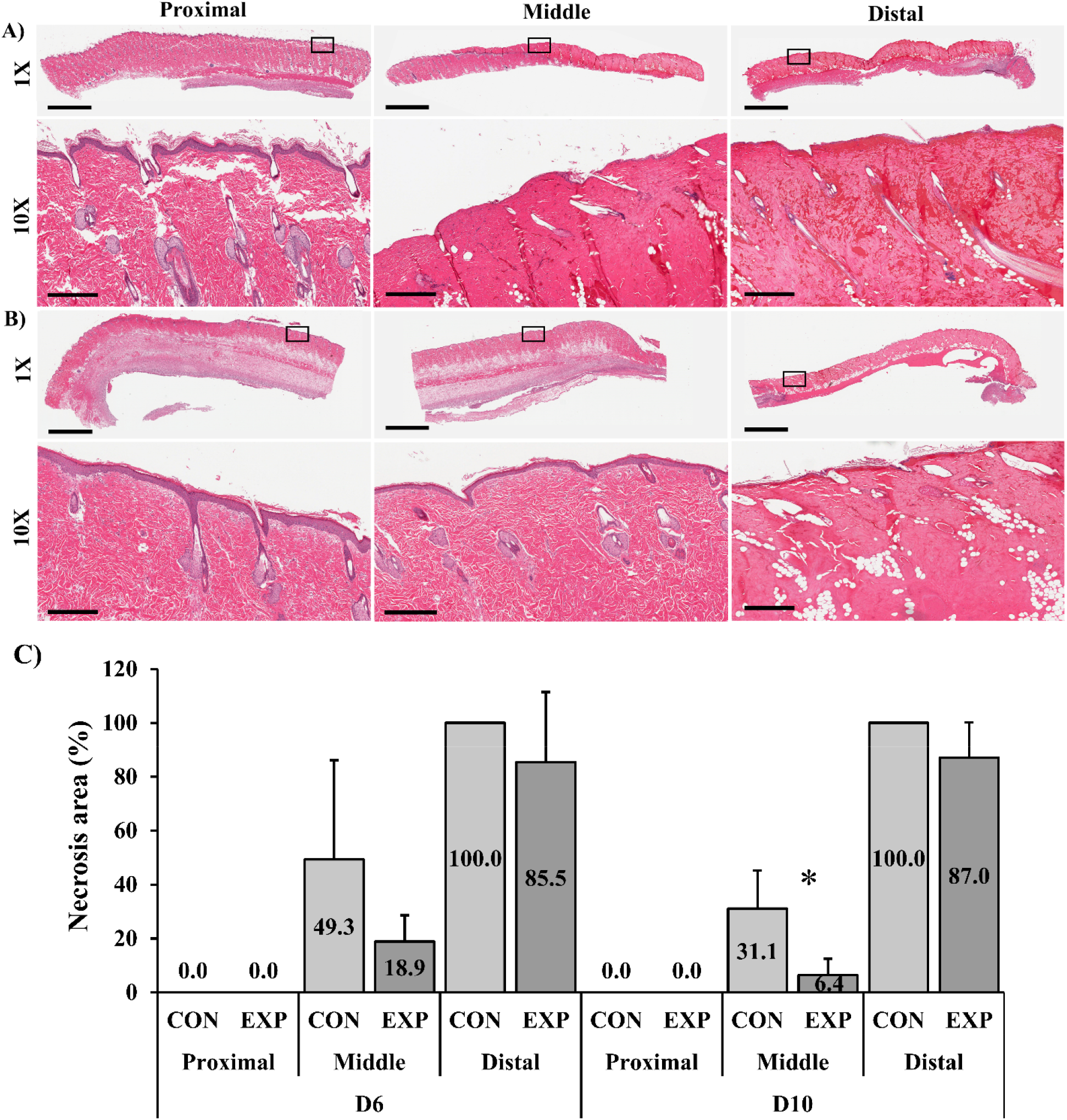
Proximal, middle, and distal area of the H&E-stained sections of skin flap in (A) control and (B) experimental group after 6 days. Black square boxes indicate the locations examined under higher magnification. The scale bar is 4mm and 400 µm in 1X and 10X magnification, respectively. (C) Histogram representing relative necrotic area observed histologically with H&E staining of the skin flap sections in control and experimental groups on day 6 and day 10, expressed as necrotic area over total area. Values are expressed as mean ± SD, N=5 per group on day 6 and N=3 on day 10, *P<0.05. Abbreviations: CON: control, EXP: experimental.

**Fig. 9.**
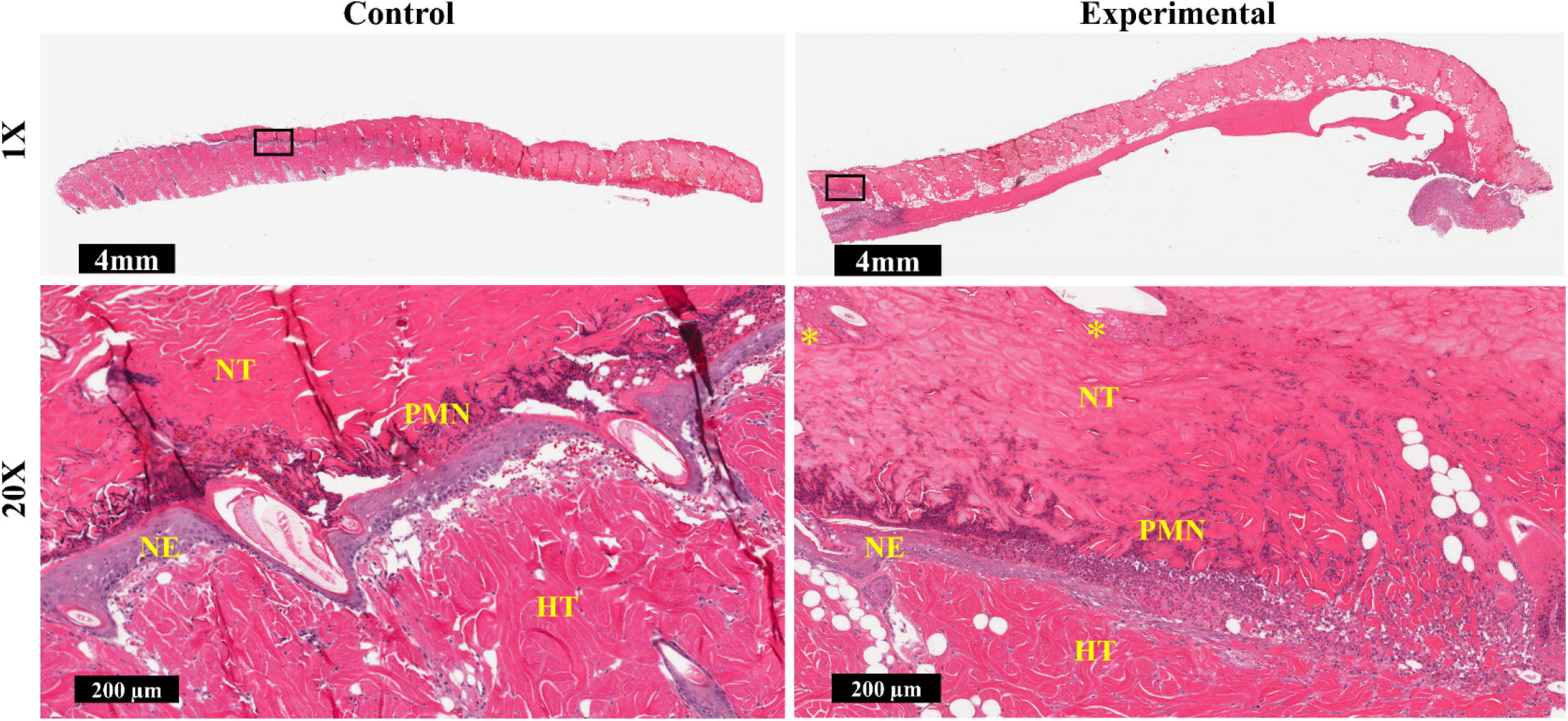
The interface of a healthy and necrotic area of H&E-stained sections of skin flap in control and experimental groups after 6 days. Black square boxes indicate the interface of healthy and necrotic tissue examined under higher magnification. HT: healthy tissue, NT: necrotic tissue, PMN: polymorphonuclear neutrophiles, NE: new epidermis and *: ghost cells.

Four days later on day 10, the necrotic area in the proximal third was 0.0% for both groups, and in the distal area we did not observe a significant difference between groups. In the mid-section, however, necrosis was significantly higher in the control group with 31.1 ± 14.1% of necrotic area versus 6.4 ± 6.1% for experimental (P<0.05).

Immunohistochemistry staining was carried out for CD34 to compare blood vessels density in control and experimental groups on day 6. Image analysis of CD34 staining (Fig. 10A, B) exhibited significantly more CD34-positive vessels between the middle and distal area of the experimental group (P<0.05), while there was no significant difference between CD34-positive vessels of middle and distal sections in the control group. Measurements of the blood vessel density (mm^2^ of blood vessels per mm^2^ of tissue) revealed no significant difference between control and experimental groups in three different sections (Fig. 10C).

**Fig. 10.**
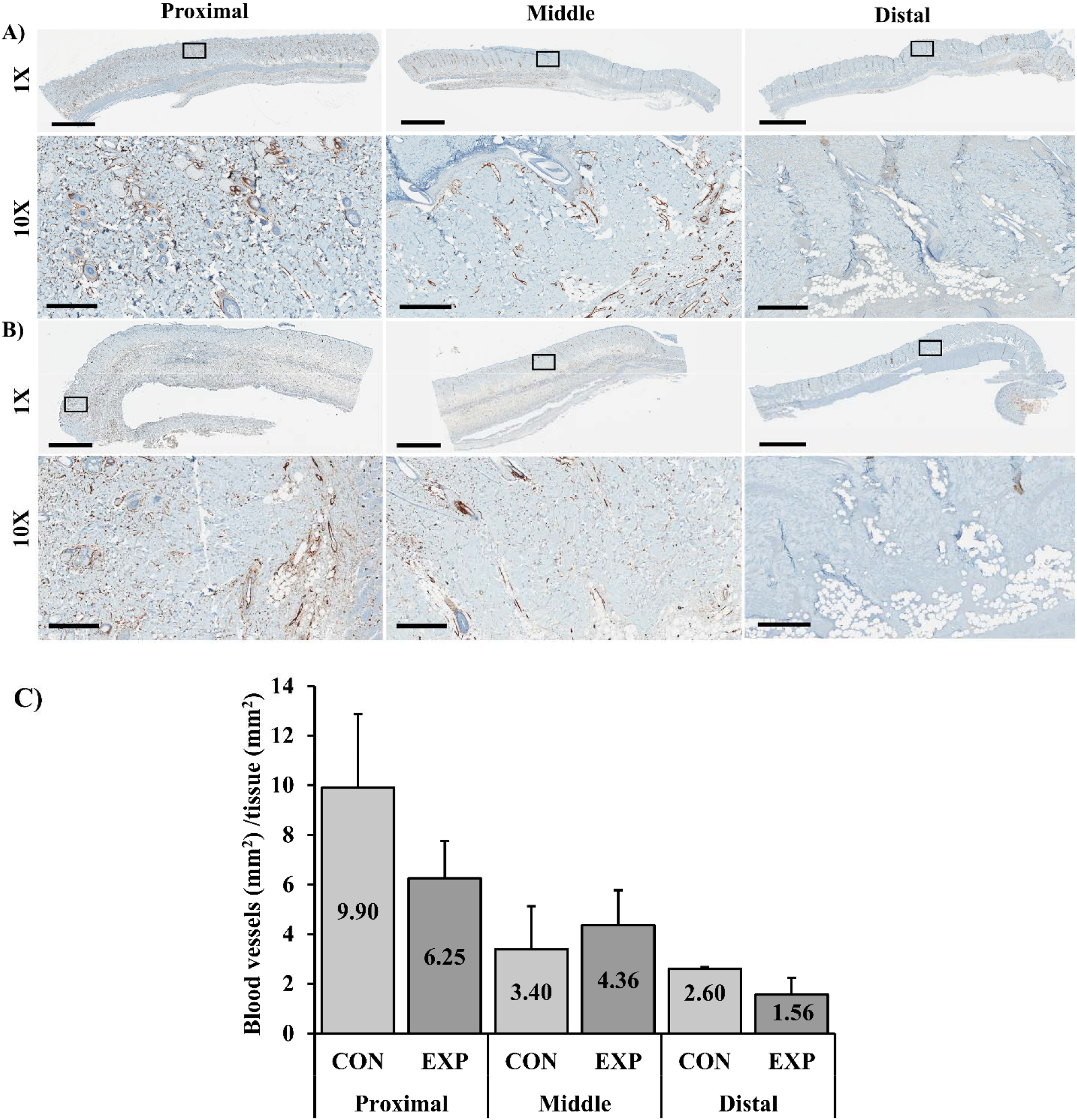
CD34-positive vessels of proximal, middle, and distal skin flap sections in (A) control (B) experimental group. The scale bar is 4mm and 400 µm in 1X and 10X magnification, respectively. (C) Histogram representing relative blood vessel density in epidermis, dermis, and hypodermis, expressed as mm^2^ of blood vessels per mm^2^ of tissue for different skin flap sections in control and experimental groups on day 6. Values are expressed as mean ± SD, N=3 per group. Abbreviations: CON: control, EXP: experimental.

To understand how macrophages participate in skin flap necrosis, we sought to identify the phenotypes of macrophages during tissue repair. Fig. 11 and Fig 12 shows immunohistochemistry staining of the middle portion of the flap, consisting of the live and dead zone on day 6. Flap sections were stained with a pan-macrophage marker (F4/80), the traditional pro-inflammatory macrophages (M1) marker (iNOS), and anti-inflammatory macrophages (M2) marker (liver arginase). Macrophages present different markers other than iNOS and liver arginase, which are not specific for the identification of M1 or M2 macrophages, and they can be detected in other cells. Therefore, we checked and confirmed the matching between expression of F4/80 (Fig.12 B) as a general macrophage marker and expression of both iNOS and liver arginase (Fig. 11B, C). Analyzing Image showed a significant difference in density for hepatic arginase with 1.36 ± 0.45/mm^2^ and 16.01 ± 8.08/mm^2^ as well as for iNOS density with 2.05± 0.87/mm^2^ and 4.98 ± 0.68/mm^2^ between control and experimental group, respectively (Fig. 11D).

**Fig. 11.**
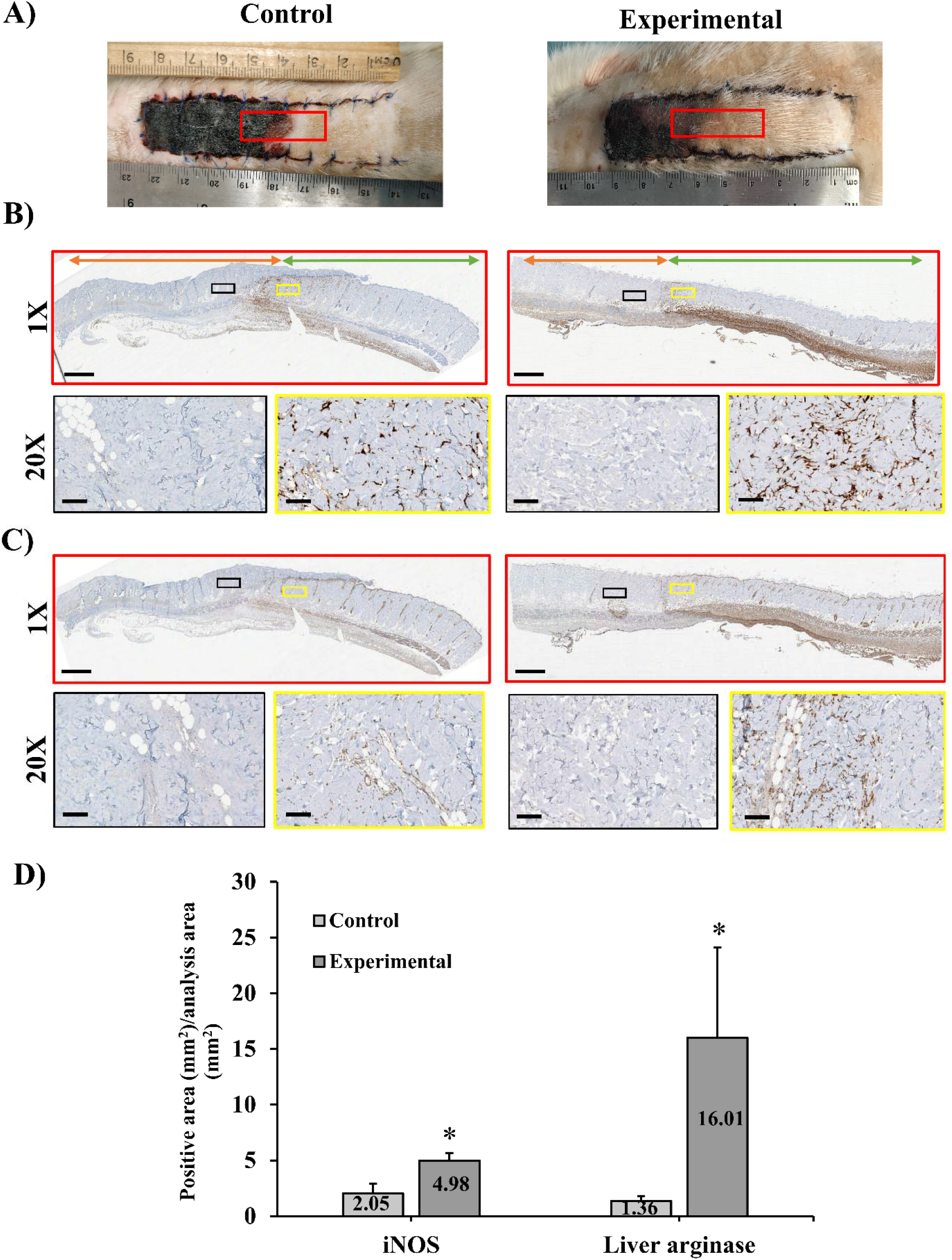
(A) Representative photographs of the skin flap on day 6 for control and experimental groups, (B) Representative iNOS and C) liver arginase immunostaining of sagittal section skin flap in middle portion for the control and experimental groups on day 6 (B) Histogram representing positive stained area (mm^2^) per analysis area (mm^2^). Values are expressed as mean ± SD, N=3 per group, *P<0.05. The scale bar is 2mm and 200 µm in 1X and 20X magnification, respectively (orange arrow: dead area, green arrow: live area).

**Fig. 12.**
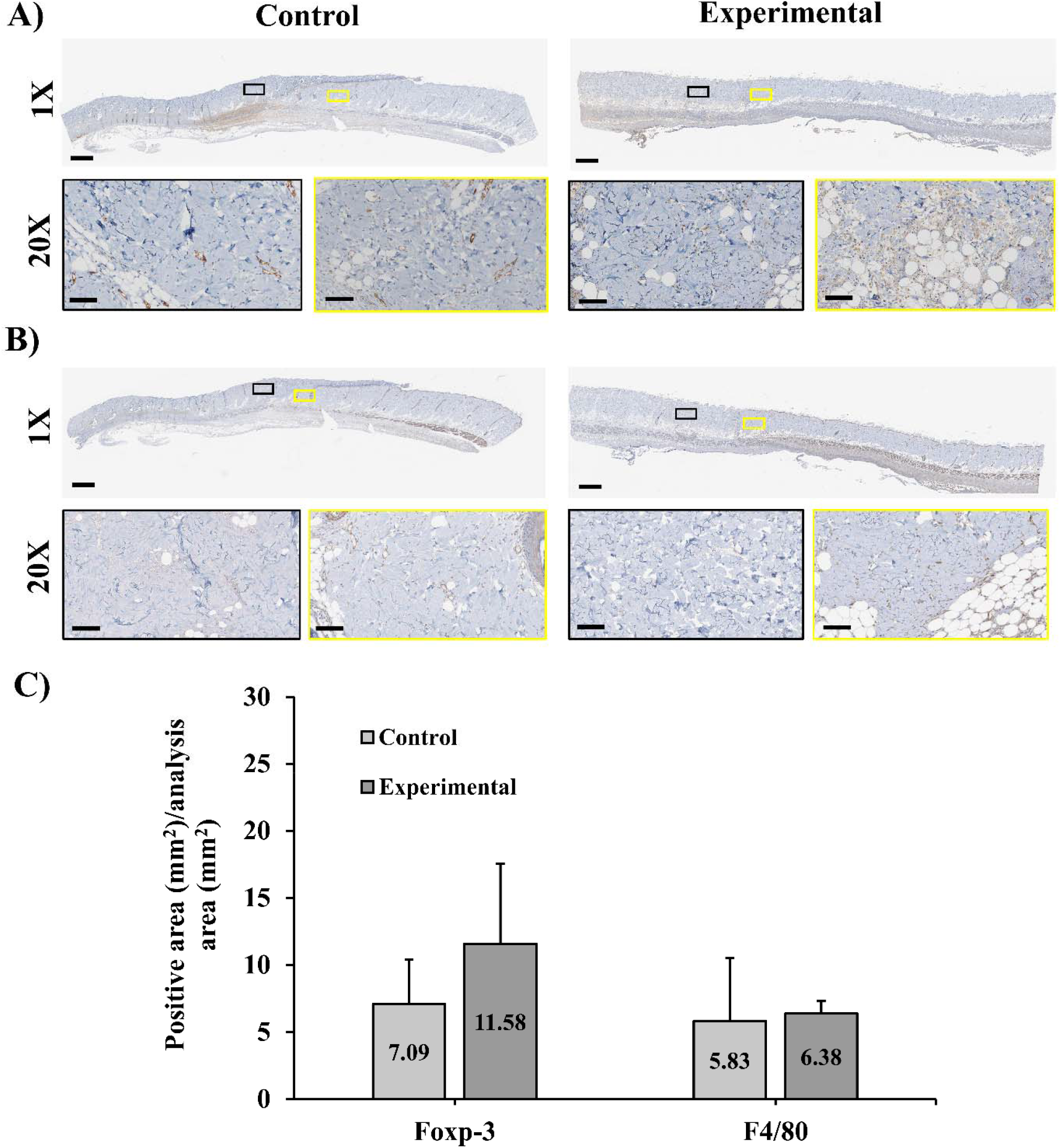
Representative (A) Foxp-3 and (B) F4/80 immunostaining of sagittal section of skin flap in middle portion for the control and experimental groups on day 6. (C) Histogram representing positive stained area (mm^2^) per analysis area (mm^2^). The scale bar is 2mm and 200 µm in 1X and 20X magnification, respectively. Values are expressed as mean ± SD, N=3 per group.

Since regulatory T cells (Tregs) play a major role in mediating immune homeostasis in the skin, we analysed Foxp3-expressing Tregs to determine whether these cells play a role in alleviating inflammation associated with wounds. We found no difference in the cell number dermis, however in the subdermal region below the necrotic viable interface a marked contrast was observed, Fig.12A, between the control and experimental group, respectively.

## 4. Discussion

Many different technologies have been developed to address issues related to oxygen insufficiency *in vitro* and *in vivo* [31], such as hyperbaric oxygen therapy (HBOT), perfluorocarbon (PFC) technologies, hemoglobin based carriers, etc. The particular advantage of the use of peroxy-compounds like calcium peroxide is the ability to produce oxygen *in situ*. Indeed to produce one liter of pure oxygen, less than ten grams, (∼3mL), of peroxide is required, while it would require several hundreds of grams of red blood cells or of PFCs or plasma to store the same volume [32]. In this study, the oxygen producing film was able to release oxygen using the reaction between calcium peroxide and water. Skin oxygen consumption is approximately 0.38µl/h/mg (∼0.017µmol/h/mg) [33], such that theoretically 2.6g of calcium peroxide could oxygenate a 7.5g skin flap for 6 days. This amount of peroxide would drastically change the subcutaneous pH and result in skin damage. Much less solid peroxide has typically been previously used preclinically. The first report by Harrison et al. [24] used oxygen generating biomaterials to prevent skin flap necrosis. In this work, sodium percarbonate was encapsulated within Poly (D, L-lactide–co–glycolide) by solvent casting method. The film was able to release oxygen over a 70h period. They showed less necrotic tissue, lower lactase concentration, and levels of apoptotic cells over 3 days. However, after 7 days, there was no significant difference between oxygen producing and control groups in the amount of tissue necrosis. Shiekh et al. [34] generated CaO_2_-encapsulated antioxidant polyurethane (PUAO) and implanted it in a skin flap on the backs of mice. A sustained oxygen release was observed from CaO_2_-PUAO for 10 days. CaO_2_-PUAO group demonstrated more skin flap survival compared to the PU scaffold without CaO_2_. Although the results confirmed less tissue necrosis through incorporating CaO_2_ into PUAO polymer compared with PU alone, no significant difference was reported in skin flap survival (histologically and visually) between no scaffold and CaO_2_-PUAO scaffold.

Here, we reduced hydrogen peroxide generation by adding a catalyst (iron oxide), a compound known to react with hydrogen peroxide (bicarbonate), and a hydrophobic polymer barrier (polycaprolactone) (Fig. 2). We also adjusted pH with an acidic buffering salt, calcium dihydrogen phosphate (Fig. 1). This reduced cytotoxicity *in vitro* since iron oxide reduced hydrogen peroxide release and calcium dihydrogen phosphate controlled shifting pH values (Fig. 3). We were able to deliver oxygen subcutaneously for 10 days (Fig. 4) and have shown significant improvement in skin flap survival over those 10 days as compared to control (Fig. 5), yet it was not sufficient to prevent necrosis in the distal portion. The results obtained for the control group in this study are consistent with other studies in literature using a similar model and a similar size [35-37].

Histomorphometric analysis of the H&E-stained sections on day 6 did not show a significant difference in total necrotic area between untreated control flaps (40.9 ± 14.1%) and oxygen treated flaps (27.3 ± 11.6%), similar to what was observed in macroscopic evaluation of necrotic areas. On day 10, the total necrotic areas were significantly different, with 35.6 ± 3.8% and 24.7 ± 5.5% for the control and oxygen generating groups, respectively (P<0.05). Previously, Bayramicli et al. [38] showed the higher contraction rate of necrosis tissue compared to the viable portion of an epigastric flap, leading to decreased necrotic area percentage at the experiment endpoint, however we corrected for contraction which occurred mainly across the width of the flap.

Another limitation of this experiment as evidenced by the skin’s blue color post-surgery, indicating the venous stasis, a pathology that commonly occurs in skin flaps and that may have participated in necrosis occurrence and spreading [39] and may have partially hindered the potential benefits of oxygen generating film. This pathology is usually addressed clinically using compression methods [40], antithrombotic [41], or leeches [42, 43]. One can easily envisage a combination of oxygen releasing materials to other techniques to extend further skin flap survival. The use of leeches has been shown to improve epigastric flap survival during venous congestion [44], and in a similar model the use of some antithrombotic has also been shown to improve skin flap viability [45]. Beyond the prevention of venous congestion, other methods have been shown to increase skin flap survival, like N-acetylcysteine [46, 47], antioxidant scavenging the radicals formed during the ischemic cascade, or vasodilators like nitroglycerine [48] and Sildenafil [49]. However, there are controversial results regarding the advantages of vasodilators. We ran a pilot to determine efficacy of these approaches and observed about 50% necrosis very rapidly after surgery by applying nitroglycerine patches topically (Fig. S1). In the absence of blood flow, it is hard to conceive how vasodilation could improve outcome [50]. Adjusting the dose, administration route, and treatment duration may have had better outcomes but further study of this approach was beyond the scope [51].

*In vivo* oxygen measurements showed that oxygen generating films could induce near physoxic oxygen concentration at any section of the flap, but the oxygen level was statistically similar between the experimental and control group at the distal portion (Fig. 6). Lack of adequate blood vessels inhibited oxygen diffusion to full skin thickness because oxygen diffusion distance through tissues is rarely more than 200µm [52], yet the thickness of the skin was several millimeters, implying that a part of the flap was not receiving oxygen from the oxygen releasing film. More blood vessels in the middle and proximal sections (higher skin temperature in thermal images) could increase oxygen levels in blood vessels and provide oxygen delivery to greater skin thickness through oxygen diffusion between vessels and cell membranes. Therefore, it is possible that the use of split thickness skin flap could result in improved viability in the distal area. The lactate concentration, a marker of tissue hypoxia, inside each section of the flap was not significantly different between both groups (Fig. 7). This suggests that oxygen delivery alone was not sufficient to maintain aerobic mechanism in the whole skin thickness.

In inflammation, wound macrophages play a key role in the skin flap due to their function in the production of mediators and growth factors at the wound site [53, 54]. Successful healing of skin flap tissue in healthy rats is characterized by the presence of alternately activated macrophages during healing [55]. M1 macrophages play a pro-inflammatory role, present antigens, and function as an immune monitor with the expression of pro-inflammatory cytokines, such as Il-6, iNOS, HLA-DR, and the transcription factors pSTAT1 and RBP -J [56-58]. M2 macrophages mainly secrete Arginase-I, interleukin 10 (IL-10), CD163, and transforming growth factor-β (TGF-β) [59-61] and other anti-inflammatory cytokines, which are responsible for reducing inflammation and play a role in wood healing [62].

Immunostaining (Fig. 11,12) showed that subdermally there was an accumulation of macrophages and that at the interface between viable dermis, (in which nuclei could be detected and collagen organisation was normal) and cell-free necrotic tissue, macrophages were present that often were found throughout the thickness of the skin. It was not possible to determine their origin, but it appeared that they migrated from below the dermis into the disorganised dermal structure, often following natural channels such as hair follicles and other appendages. The expression of iNOS and liver arginase positive cells at least partially matched the localisation of expression of pan-macrophage marker F4/80. Macrophage M2 phenotype is associated with the promotion of tissue repair and pro-angiogenic function [63, 64], suppression of inflammation (by limiting responses dependent on M1), and regulation of wound healing and fibrosis [65].

After the skin injury the inflammatory phase occurs early, and it is characterized by a strong activation of the innate and adaptive immune systems. Tregs constitute a large percentage of lymphocytes that reside in the skin [66, 67], and they play an important role in the regulation of tissue inflammation. We sought to determine whether Tregs play a role in the healing of skin wounds in our experimental group compared to control. We found that there was a slight, non-significant increase in the subdermal tissue in rats with oxygen releasing film compared to control. Knowing that activated Tregs accumulate in the skin and play a major role in reducing the accumulation of pro-inflammatory macrophages, which it does explain the low rate of M1 in control compared to the experimental. Maybe the oxygen releasing film increased the number of M1 cells to increase the transition to M2.

It was not possible to determine if the difference in polarisation was caused by the oxygen treatment or were simply an effect of the improved healing, although the possibility of immunomodulation in disruption of necrosis is under active study by other groups [68-71]. However, previously increased polarization of M2 macrophages has been observed to coincide with the oxygen level in the microenvironment [72, 73]. The sequential occurrence of the two macrophage polarization states has been reported as being necessary for correct resolution of healing [63, 74] and for tissue repair after injury [75]. Given the complexity of the healing, inflammatory, and host bed response, it appears not unreasonable that elevating oxygen levels in what is otherwise ischemic tissue may have multiple beneficial effects to mitigate conditions that are conducive to ischemic necrosis.

## 5. Conclusion

Here we report the fabrication and use of a non-biodegradable oxygen delivery implant which was able to sustain the release of a large amount of oxygen over 14 days. Although we demonstrate its efficacy to prevent necrosis, the study is inherently limited by the oxygen amount delivered and by the ischemic model used. Oxygen is not the only factor of necrosis onset and venous congestion that may have played a non-negligible role, nonetheless oxygen delivery may have an adjuvant role with other conventional therapies to further prevent necrosis onset.

## Data Availability

The raw/processed data required to reproduce these findings cannot be shared at this time as the data also forms part of an ongoing study. Sections of data may be available on request.

## Supplementary Data

**Fig. S1.**
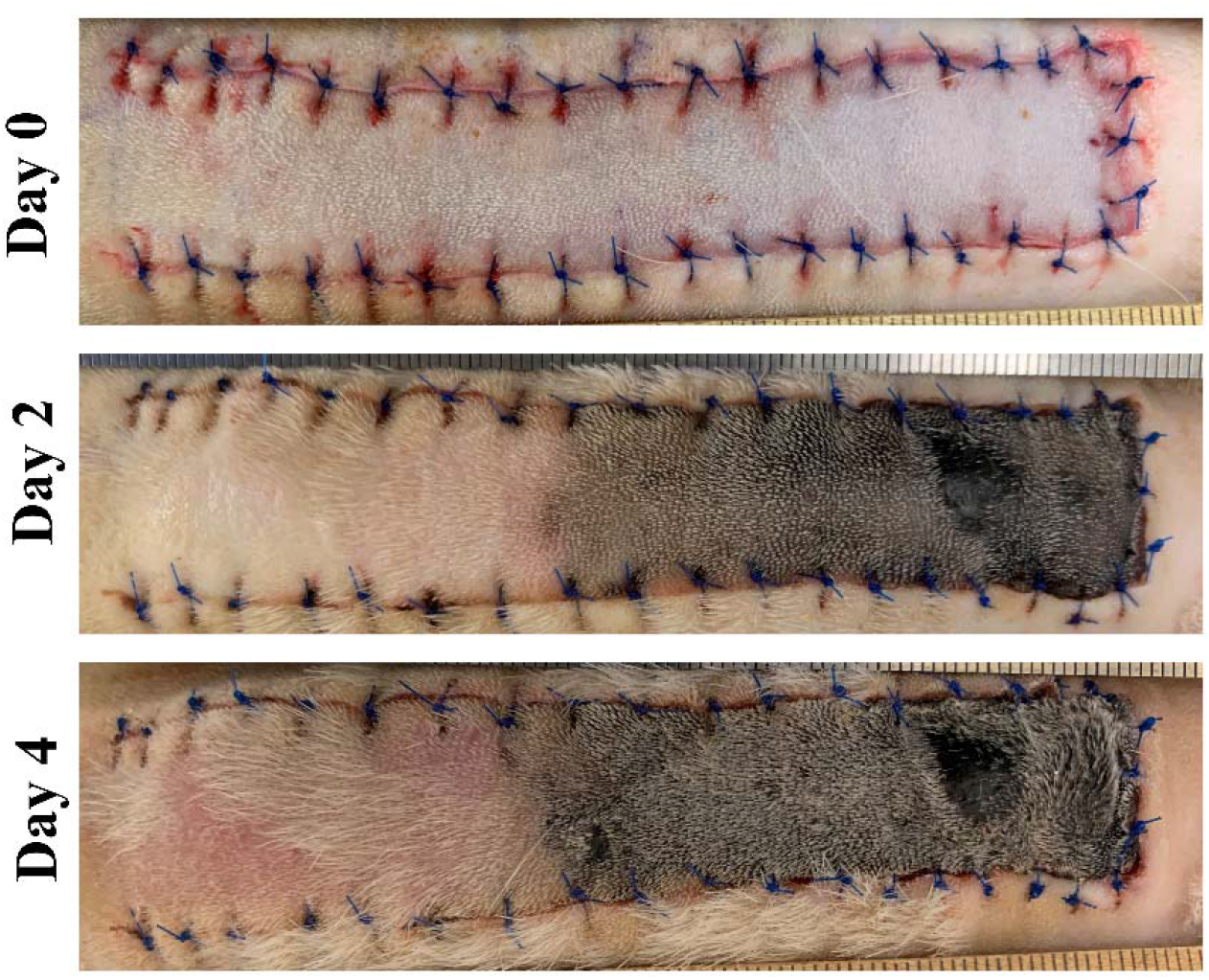
Representative photographs of the skin flap at days 0, 2 and 4 for nitroglycerine. Two Trinipatch 0.2 mg/hr, 7cm^2^ were applied after the operation and days 1, 2 and 4 over the flap

